# Divergent SARS-CoV-2-specific T and B cell responses in severe but not mild COVID-19

**DOI:** 10.1101/2020.06.18.159202

**Authors:** Anna E. Oja, Anno Saris, Cherien A. Ghandour, Natasja A.M. Kragten, Boris M. Hogema, Esther J. Nossent, Leo M.A. Heunks, Susan Cuvalay, Ed Slot, Francis H. Swaneveld, Hans Vrielink, Theo Rispens, Ellen van der Schoot, René A.W. van Lier, Anja Ten Brinke, Pleun Hombrink

**Affiliations:** Department of Hematopoiesis, Sanquin Research and Landsteiner Laboratory, Amsterdam UMC, University of Amsterdam, Amsterdam, the Netherlands; Centre for Experimental and Molecular Medicine, Amsterdam UMC, Amsterdam, the Netherlands; Amsterdam UMC Covid study group, Amsterdam UMC, Amsterdam, the Netherlands; Sanquin Diagnostic Services and Sanquin Research, Amsterdam UMC, University of Amsterdam, Amsterdam, the Netherlands; Department of Pulmonary Medicine, Amsterdam UMC, Amsterdam, the Netherlands; Department of Intensive Care Medicine, Amsterdam UMC, Amsterdam, the Netherlands; Unit of Transfusion Medicine, Sanquin Blood Supply, Amsterdam, the Netherlands; Laboratory of Blood-borne Infections, Sanquin Blood Supply, Amsterdam, the Netherlands; Department of Immunopathology, Sanquin Research and Landsteiner Laboratory, Amsterdam UMC, University of Amsterdam, Amsterdam, the Netherlands; Department of Experimental Immunohematology, Sanquin Research and Landsteiner Laboratory, Amsterdam UMC, University of Amsterdam, Amsterdam, the Netherlands

## Abstract

Severe acute respiratory syndrome coronavirus 2 (SARS-CoV-2) is the causative agent of the current coronavirus disease 2019 (COVID-19) pandemic. Understanding both the immunological processes providing specific immunity and potential immunopathology underlying the pathogenesis of this disease may provide valuable insights for potential therapeutic interventions. Here, we quantified SARS-CoV-2 specific immune responses in patients with different clinical courses. Compared to individuals with a mild clinical presentation, CD4+ T cell responses were qualitatively impaired in critically ill patients. Strikingly, however, in these patients the specific IgG antibody response was remarkably strong. The observed disparate T and B cell responses could be indicative of a deregulated immune response in critically ill COVID-19 patients.

---

The novel severe acute respiratory syndrome coronavirus 2 (SARS-CoV-2) is a single-strand RNA virus that has been identified as the causative agent of coronavirus disease 2019 (COVID-19), mainly characterized by respiratory symptoms. This virus appears to be easily transmittable and highly virulent in a sizable fraction of the global population. Consequently, the WHO characterized COVID-19 as a pandemic by March 11^th^, 2020. Despite socioeconomic lockdown in about a third of the world, the global number of confirmed SARS-CoV-2 infections surpassed 7 million^1,2^(https://covid19.who.int/).

SARS-CoV-2 is primarily transmitted via respiratory droplets and aerosols. The virus infects airway epithelial cells expressing the surface receptors ACE2 and TMPRSS2. The virus replicates and sheds virus particles, which cause the infected cell to undergo pyroptosis. During this process, damage-associated molecular patterns are released, activating a cascade of pro-inflammatory cytokines and chemokines that attract auxiliary immune cells to the site of infection promoting further inflammation^3^. In severely affected COVID-19 patients this pro-inflammatory cascade appears to be dysregulated, resulting in a life-threatening cytokine storm, which triggers acute respiratory distress syndrome (ARDS) and subsequently other organ dysfunction including, liver, heart, and kidney damage.

Clinically, COVID-19 patients can be classified into mild, moderate, severe, and critical cases. Importantly, it has been proposed that the immune response against SARS-CoV-2 is linked to the severity of disease^2,4,5^. Although T cell responses against SARS-CoV-2 proteins have been characterized^6,7^, a comprehensive comparison of the quantity and quality of adaptive immune responses in patients with distinct clinical courses is yet to be performed. Among the cells recruited to the lungs are virus-specific T cells, that have been primed by dendritic cells in the lung draining lymph nodes, and set to kill virus-infected cells, thus preventing further spread and thereby limiting disease progression. Both SARS-CoV-2-specific CD8^+^ and CD4^+^ T cells and T cells with an activated phenotype have been detected in the blood between 1 and 2 weeks after the onset of COVID-19 symptoms^6,8,9^. These SARS-CoV-2-specific T cells predominantly produced Th1 cytokines, although Th2 cytokines were also detected, especially in critically ill patients. CD8^+^ T cells are important for directly attacking and killing virus-infected cells, whereas CD4^+^ T cells also prime B cells. In particular, a subset of CD4+ T cells, follicular helper T cells (Tfh) are crucial for the survival and class-switching of B cells. Tfh frequencies increase in blood over the course of SARS-CoV-2 infection and are increased in critically ill patients compared to healthy individuals^7,8^. Intriguingly, many patients with severe COVID-19 develop a global T cell lymphopenia, which may reflect massive recruitment to the lungs and other inflamed tissues^2,5,10^. An alternative mechanism that could contribute to the disappearance of circulating T cells may be high levels of activation-induced cell death of lymphocytes in the secondary lymphoid organs^11^.

The accumulation of exhausted T cells with a reduced functional diversity in peripheral blood of COVID-19 patients with severe symptoms further supports the notion that antigen-specific T cells are instrumental in the control of SARS-CoV-2 infection^12,13^.

We studied the characteristics of SARS-CoV-2-specific adaptive immune responses in COVID-19 individuals with distinct symptoms, ranging from mild and severe to critically ill patients. This type of comparison is essential to understanding the contribution of T and B cell responses to viral clearance and the mechanisms behind immunopathology. We observe major differences in the quality and quantity of these responses across the different clinical groups. This information is not only of eminent relevance for understanding the pathophysiology of COVID-19, but also could be helpful in diagnosis, treatment, and vaccine design.

## Results

### T cell activation and lymphopenia are associated with disease severity in COVID-19

Peripheral blood of a total of 56 PCR-positive COVID-19 patients was collected, of whom 37% (n=21) recovered from disease and expressed mild symptoms (mild symptoms up to mild pneumonia), 25% (n=14) recovered from disease and expressed severe symptoms requiring hospitalization and 38% (n=21) critically ill requiring treatment in intensive care unit (ICU), referred to as mild, severe, and ICU groups from now on, respectively (Table 1, Extended Data Table 1). For all patients SARS-CoV-2 exposure was verified by serology testing of IgG titers against RBD of spike and nucleocapsid. For the ICU cohort, blood was taken during ICU stay for 18 patients, indicated in figures by square symbols, and after recovery for 3 patients, shown as circles in all figures. Samples were collected with at a median of 52 (mild), 45 (severe) and 34 (ICU) days after onset of symptoms (Table 1, Extended Data Table 1).

**Table 1:**

|  | Bloodbank donors | Not-hospitalized | Hospitalized | ICU |
| --- | --- | --- | --- | --- |
| Severity | Unexposed | Mild* | Severe* | Critical* |
| Number | 16 | 21 | 14 | 21 (3<sup>#</sup>) |
| Age (yr) | 52 [26-67] | 43 [23-63] | 52 [30-61] | 63 [32-77] |
| Gender (F/M) | 8 [50%] / 8 [50%] | 8 [38%] / 13 [62%] | 3 [21%] / 11 [79%] | 2 [10%] / 19 [90%] |
| Sampling (day)** | N/A | 52 [25-72] | 45 [21-67] | 34 [11-59] |

\*SARS-CoV-2 PCR positive
\*\*Days after onset of symptoms
<sup>#</sup>ICU recovered at sampling time

First the frequency of T cells in the blood of different COVID-19 patient groups was assessed by fluorescence-activated cell sorting (FACS) (general gating strategy shown in Extended Data Fig.1a). The proportion of CD3+ T cells was decreased in the critical patient group during ICU stay as compared to healthy controls and T cell levels remained low for the severe patients at a median of 30 days after symptomatic recovery (Fig. 1a, Extended Data Fig. 1b). The decreased peripheral T cell frequencies were observed equally in CD4+ and CD8+ cells without a change in CD4/CD8 ratio (Fig. 1b, Extended Data Fig. 1c). The severe and ICU patient groups demonstrated higher frequencies of effector phenotype (CD45RA+CD27-) and severe patients had lower frequencies of naïve (CD45RA+CD27+) CD4+ T cells than mild patients or unexposed individuals (Fig. 1c). Effector CD8+ T cells were also relatively increased in the ICU patients compared to patients with mild or severe symptoms, while naïve CD8+ T cells were significantly decreased compared to unexposed individuals and mild patients (Fig. 1d). However, we cannot exclude that the skewed effector phenotype in the ICU patients is not due age or CMV seropositivity.

**Fig. 1:**
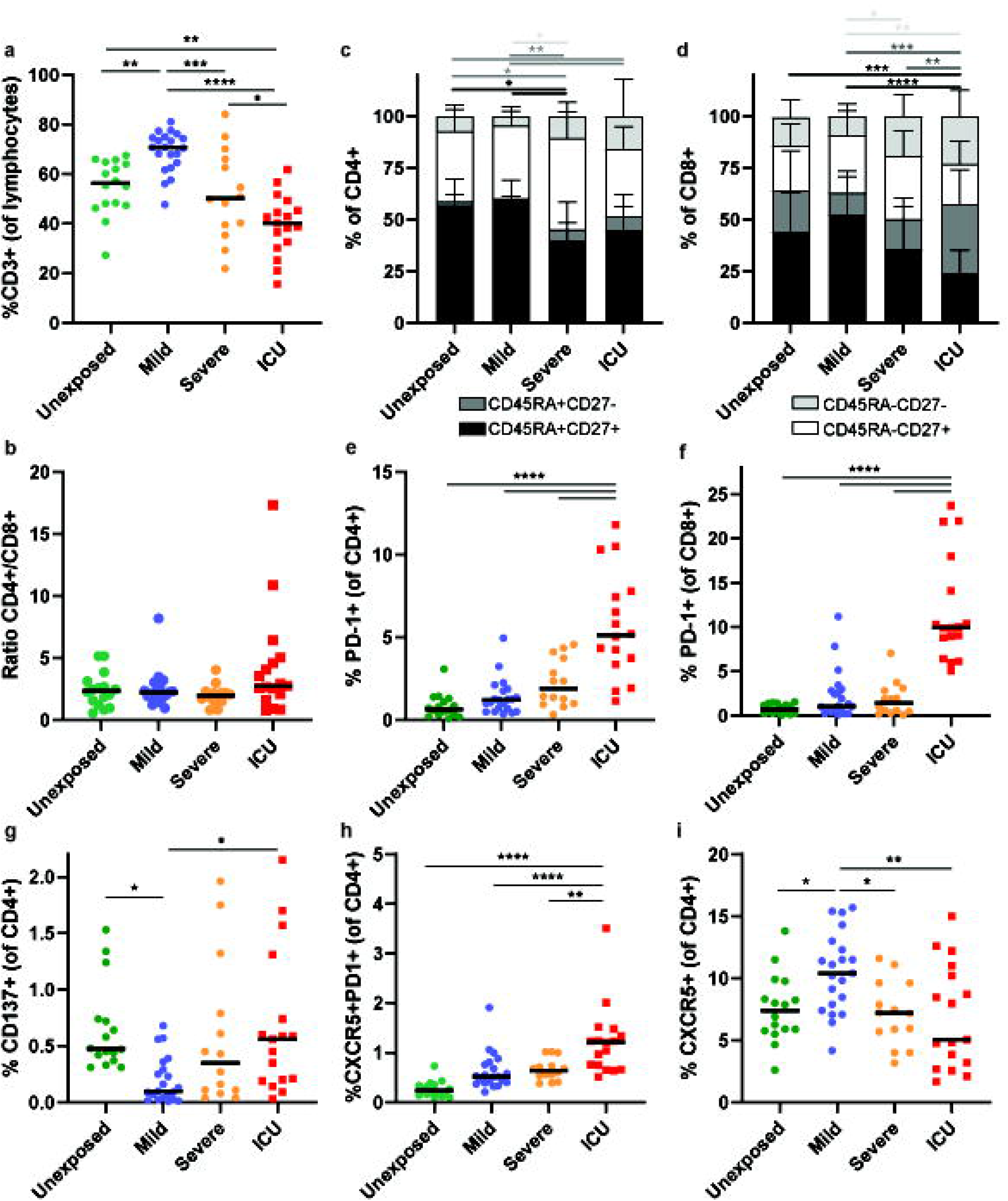
T cell activation and lymphopenia are associated with Covid-19 disease severity. **a,** Overview of donor/patient characteristics used in this study. **b,c**, Frequency of CD3+ T cells of total lymphocytes and the ratio of CD4+ to CD8+ T cells was determined in unexposed individuals (green), and Covid-19 patients with mild (blue), severe (orange), or critical (ICU) (red) symptoms, symbol colors are the same for all figures. Circles indicate samples obtained after clearance of symptoms during recovery phase. Squares indicate samples obtained during ICU stay. **d,e**, The composition of CD4+ and CD8+ T cells was determined based on CD45RA and CD27 expression, to identify naïve (black; CD45RA+CD27+), effector (dark grey; CD45RA+CD27-), memory (white; CD45RA-CD27+), and effector memory (light grey; CD45RA-CD27-) cells. **f-j**, Frequency of PD-1+ of CD4+, PD-1+ of CD8+, CD137+CD40L- of CD4+, cTfh (CXCR5+PD1+) of CD4+ and CXCR5+ of CD4+ T cells was determined in the different cohorts. **b,c,f-j**, Each symbol indicates a unique sample and the line indicates the median values. Circular symbols depict donors that recovered from Covid-19 and square symbols depict patients in the ICU at the time of sampling throughout the manuscript. Significance of the results was determined using one-way ANOVA with Tukey’s multiple comparisons test. **d,e**, Bars show mean values with SD and the significance was determined using two-way ANOVA with Tukey’s multiple comparisons test. N=16, N=21, N=14, and N=17 donors/patients for unexposed, mild, severe, and ICU cohorts respectively.

We next analyzed the cell surface expression of the inhibitory PD-1, expressed after T cell activation, to obtain a first view on the functionality of the T cell compartment. Frequencies of PD-1+ CD4+ and CD8+ T cells was increased in association with stage of infection, with the highest levels in ICU patients (Fig. 1e,f, Extended Data Fig. 1d,e). Recovered ICU patients showed significantly lower PD-1 expression than the patients in the ICU at the time of sampling (Extended Data Fig. 1i,j). Using constitutive expression of the 4-1BB receptor protein CD137 and absence of the conventional CD4+ T cell activation marker CD40L as a proxy for activated regulatory T cells (Treg)^14^, there was a significant decrease in activated Treg in blood of the mild group compared to unexposed individuals and ICU patients (Fig. 1g, Extended Data Fig. 1f,k). This may indicate recruitment of Treg to tissues in the mild cohort which can prevent immunopathologies due to hyper-inflammatory responses, frequently observed in critically ill COVID-19 patients.

It was recently shown that circulating follicular helper T cells (cTfh) are increased in Covid-19 patients and rise over the course of the infection^8,15,16^. However, possible differences in relation to disease severity were not assessed. Thus, we also analyzed the frequencies of circulating follicular T helper cells (cTfh). The highest frequencies of cTfh, based on PD-1 and CXCR5 expression, were found in ICU patients (Fig. 1h, Extended Data Fig. 1g,l). Since the recovered ICU patients showed a trend towards lower frequencies of cTfh (Extended Data Fig. 1g), it is unclear whether frequencies of cTfh correlate with stage of disease or disease severity. On the other hand, concomitant with the lymphopenia, we found fewer CXCR5 expressing CD4+ T cells in the ICU but not in the recovered mild and severe COVID-19 patient groups (Fig. i, Extended Data Fig. 1h,m). This might reflect homing to CXCL13 gradients in secondary lymphoid tissues at the site of antigen accumulation under conditions of inflammation.

### The bronchoalvelor lavage fluid of ICU COVID-19 patients contains effector-memory but few tissue-resident T cells

Following a primary respiratory viruses infection, a population of tissue resident memory T cells (T_RM_) is generated, that persist in the tissue for prolonged periods of time^17,18^. In healthy human lungs, T_RM_ express canonical phenotypic markers for retention, adhesion and migration to tissues (CD69, CD103, CD49a, CXCR6) and express high levels of the inhibitory molecule PD-1^19,20^. Since both airway CD4+ and CD8+ T cells were reported to be required for optimal protection against the emerging coronavirus SARS-CoV-2^21,22^, we set out to sample bronchoalveolar lavage fluid (BALF) and blood of 8 critical COVID-19 patients during ICU stay, for monitoring T cell phenotypes. Under healthy circumstances, T cells in BALF reflect the composition of the barrier tissue and exhibit predominant T_RM_ phenotypes^23^. The CD4:CD8 T cell ratio in BALF was not significantly different when compared to that derived from paired blood (Fig. 2a). The BALF contained lower frequencies of naïve (CD45RA+CD27+) and higher frequencies of effector memory (CD45RA-CD27-) CD4+ and CD8+ T cells compared to blood-derived PBMCs (Figure 2b,c). Nonetheless, the high frequency of naïve cells in the BALF was striking, since under normal conditions very few to no naïve cells are found in the BALF^23^.

**Fig. 2:**
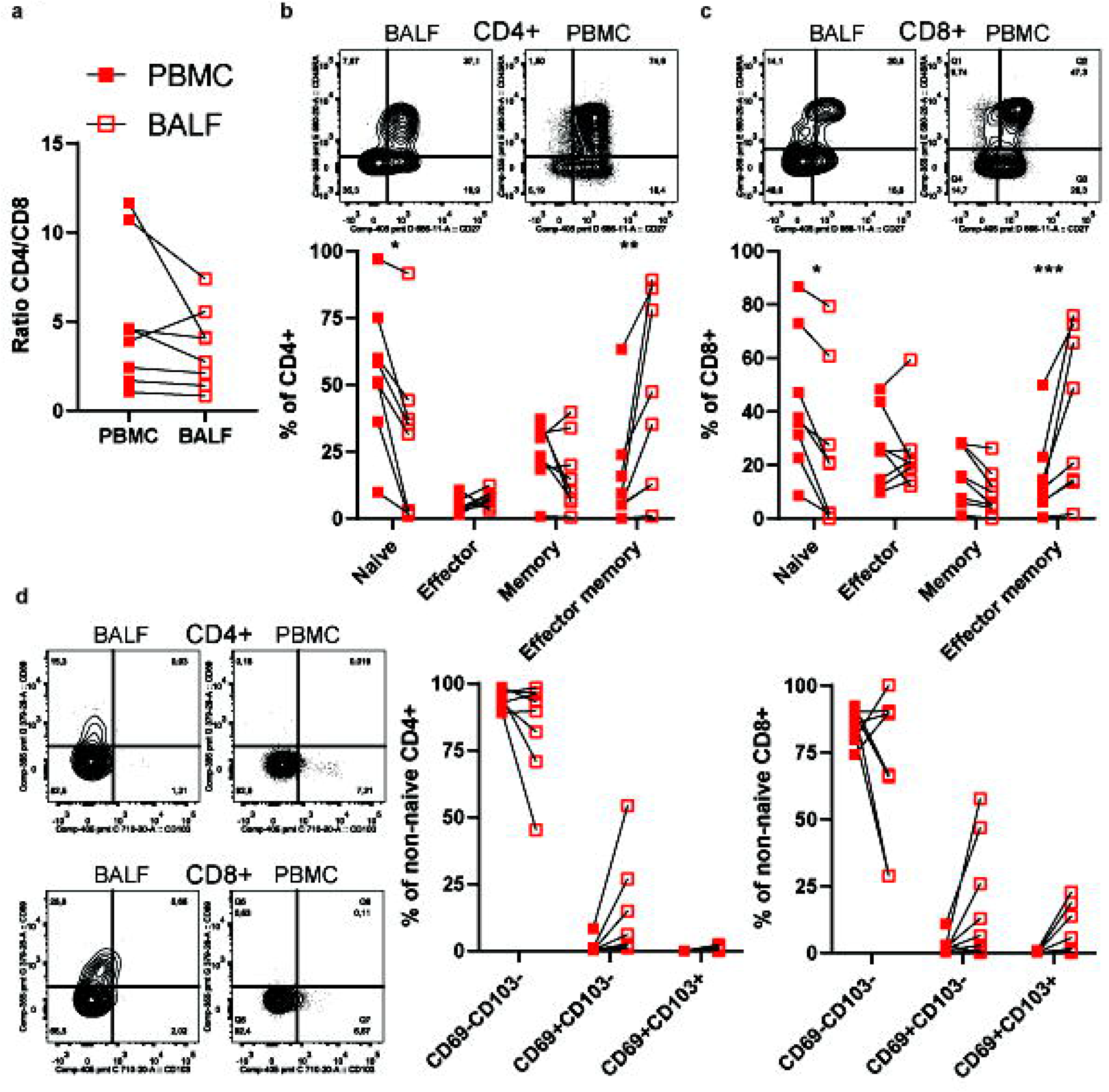
The bronchoalvelor lavage fluid (BALF) of ICU COVID-19 patients contains effector-memory but few tissue-resident T cells. **a**, Ratio of CD4+ to CD8+ T cells in paired BALF vs PBMC samples. **b,c**, The composition of CD4+ and CD8+ T cells was determined based on CD45RA and CD27 expression, to identify naïve (CD45RA+CD27+), effector (CD45RA+CD27-), memory (CD45RA-CD27+), and effector memory (CD45RA-CD27-) cells. **d,e**, Frequencies of resident memory CD4+ and CD8+ T cells (T_RM_), measured by expression of CD69 and CD103 on antigen-experience non-naïve T cells, where T_RM_ are defined as CD69+CD103- or CD69+CD103+. **a-e**, Each symbol indicates a unique sample and paired BALF and PBMC samples are connected by a line. Significance of the results was determined using one-way ANOVA with Tukey’s multiple comparisons test. **b-e**, Representative contour plots are shown for each group. N=8 patients.

We further examined whether memory T cells in the BALF exhibited a T_RM_ phenotype, by measuring CD69 and CD103 expression on non-naïve CD4+ and CD8+ T cells. In line with the abundance of T cells with a circulating phenotype, we also found very few CD4+ or CD8+ T cells expressing CD69 or CD69 and CD103 (Fig. 2d). Thus BALF T cells in ICU COVID-19 patients are not cells that have the canonical T_RM_ phenotype, but rather appear to be cells that normally belong to the circulating pool.

### Mild COVID-19 patients have broad CD4+ responses to SARS-CoV-2 peptides but low frequencies of cross-reactive T cells

To identify the immunodominant SARS-CoV-2 antigens recognized by CD4+ T cells, we stimulated PBMCs from convalescent non-hospitalized COVID-19 patients with a range of SARS-CoV-2 peptide pools and measured the upregulation of activation markers CD40 ligand (CD40L/CD154) and CD137 (4-1BB) by flow cytometry (Fig 3a, Extended data Fig 2a). Using 15-mer peptide pools, we focused the analysis on CD4+ T cells. In accordance with recent publications^6,7^, Spike (S) was the most immunodominant with a 100% response rate, followed by Nucleocapsid (N) and membrane glycoprotein (M) which were also recognized in the majority of patients. Slightly less common, ORF3A (AP3A) and a pool of open reading frame, non-structural and uncharacterized antigens (ORF pool) were also recognized by a portion of CD4+ T cells, whereas the envelope protein (E) was not (Fig. 3b). We further added up the responses to all these antigens to estimate the total CD4+ T cell response to SARS-CoV-2 and compared this to the frequencies of CMV pp65 specific CD4+ T cells. The magnitude of SARS-CoV-2 specific CD4+ T cells was significantly higher than that of CMV pp65 specific CD4+ T cells (Fig. 3c). The CMV responses varied, likely due to seropositivity, while all patients showed a robust response to SARS-CoV-2.

**Fig. 3:**
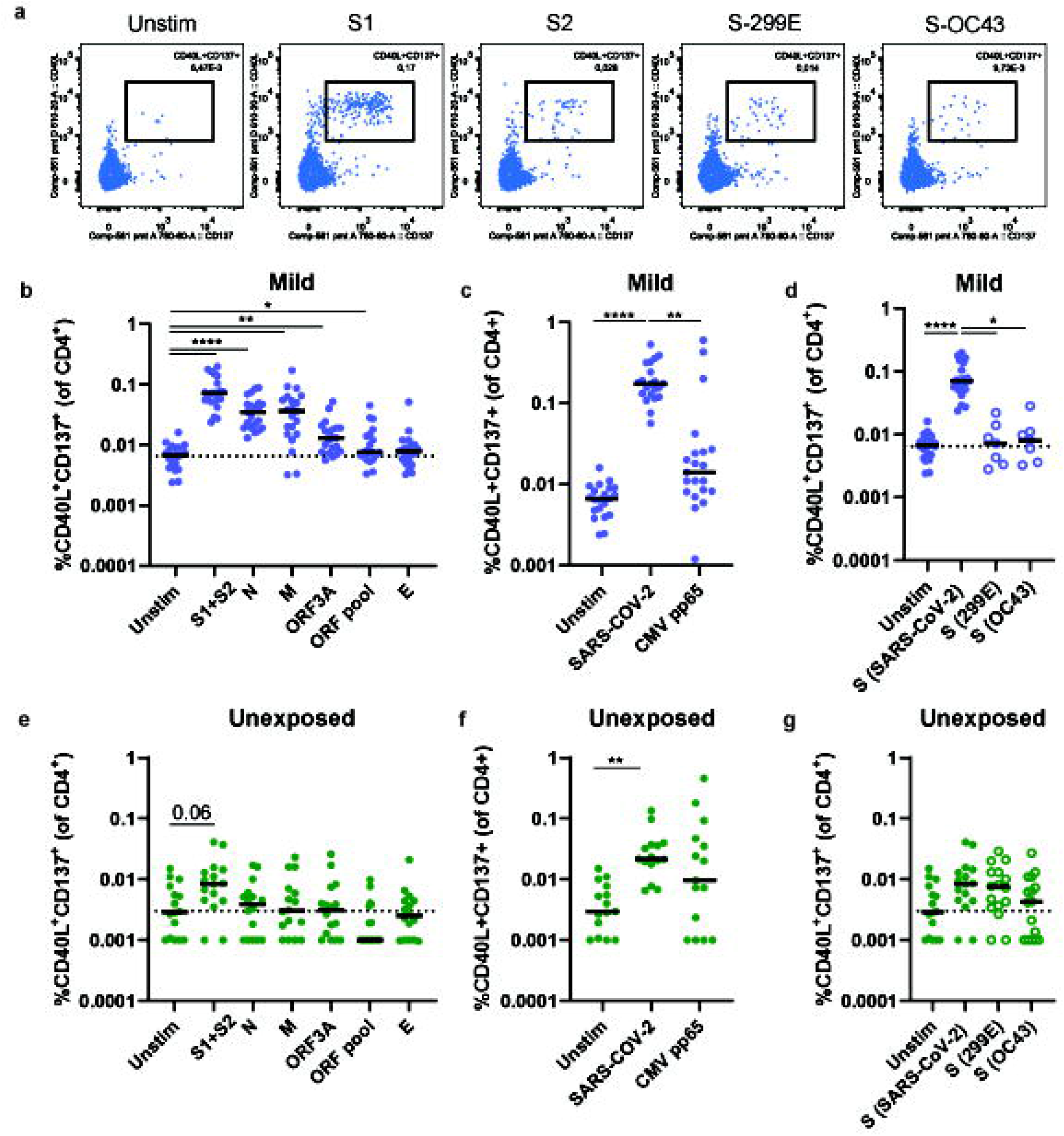
Mild COVID-19 patients have broad CD4+ responses to SARS-CoV-2 peptides but low frequencies of cross-reactive T cells. **a** Representative FACS plot examples of total CD4 T cells of a mild covid-19 patient unstimulated or stimulated with SARS-CoV-2-Spike (S) S1 and S2 pools, common-cold corona virus 229E-S and OC43-S peptide pools. **b,e** Frequencies of CD4+ T cells specific for different SARS-CoV-2 antigens, S1 and S2, Nucleocapsid (N), Membrane (M), ORF3A, ORF pool and Envelope (E), was determined by the upregulation of CD40L and CD137 upon peptide stimulation and compared to the unstimulated (Unstim) in mild Covid-19 patients (**b**) and unexposed donors (**e**). **c,f**, The responses to the different antigens was added up to calculate the total SARS-CoV-2 response and compared to frequencies of CMV-pp65 specific CD4+ T cells in mild Covid-19 patients (**c**) and unexposed individuals (**f**). **d,g**, Cross-reactivity against spike antigen from other coronavirus strains was assessed by comparing SARS-CoV-2-S CD4+ responses to 299E-S, and OC43-S CD4+ responses in mild Covid-19 patients (**d**) and unexposed individuals (**g**). **b-g**, Each symbol indicates a unique sample and the line indicates the median values. Significance of the results was determined using one-way ANOVA with Tukey’s multiple comparisons test. N=16 and N=21 for unexposed and mild cohorts respectively.

To investigate whether these high responses against S were SARS-CoV-2-specific or if they also consisted of cross-reactive T cells to “common cold” coronavirus strains, we stimulated these same PBMC samples with two Spike peptide pools, 299E and OC43, from two different “common cold” coronavirus strains. The frequencies of SARS-CoV-2 S-specific CD4+ T cells were significantly higher than S-299E- or S-OC43-specific CD4+ T cells (Fig. 3d). Furthermore, the frequencies of 299E- and OC43-specific CD4+ T cells were not significantly higher than the paired unstimulated samples. This demonstrates that the responses we observe against S were predominantly due to COVID-19.

In line with this, we also determined the responses of unexposed healthy donor PBMCs to these SARS-CoV-2 antigens (Extended data Fig 2b). While the response frequencies to the individual antigens were not significantly higher than the unstimulated, there was a trend towards higher frequencies of S-specific CD4+ T cells (Fig. 3e). When the low frequent responses were added up this was higher than the unstimulated, but not significantly different from frequencies of CMV pp65 specific CD4+ T cells (Fig. 3f). We further compared the CD4+ T cell responses to SARS-CoV-2-S and “common cold” S-299E and S-OC43 peptide pools and found no differences in the frequencies between the three spike peptide pools (Fig. 3g). These data indicate that unexposed healthy donors have low frequencies of cross-reactive CD4+ T cells to S of SARS-CoV-2.

### COVID-19 in the ICU patients have diminished frequencies of SARS-CoV-2-specific CD4+ T cells that are also functionally impaired

After identifying S, N, and M as the immunodominant antigens of SARS-CoV-2, we assessed the CD4+ T cell responses in the two additional patient cohorts, severe (hospitalized) and ICU (Extended Data Fig 2c,d). Also in these patient cohorts, S was most frequently recognized, followed by N and M (Fig. 4a,b). The majority of the S- and N-specific CD4+ T cells had a memory (CD45RA-CD27+) phenotype. (Fig. 4c,d). We also sought to determine the antigen-specificity of CD4+ T cells in paired BALF and PBMC samples. In one sample, we found S-specific CD4+ T cells in both BALF and PBMCs and in both tissues the S-specific CD4+ T cells had a circulating memory rather than a T_RM_ phenotype (Extended Data Fig. 3).

**Fig. 4:**
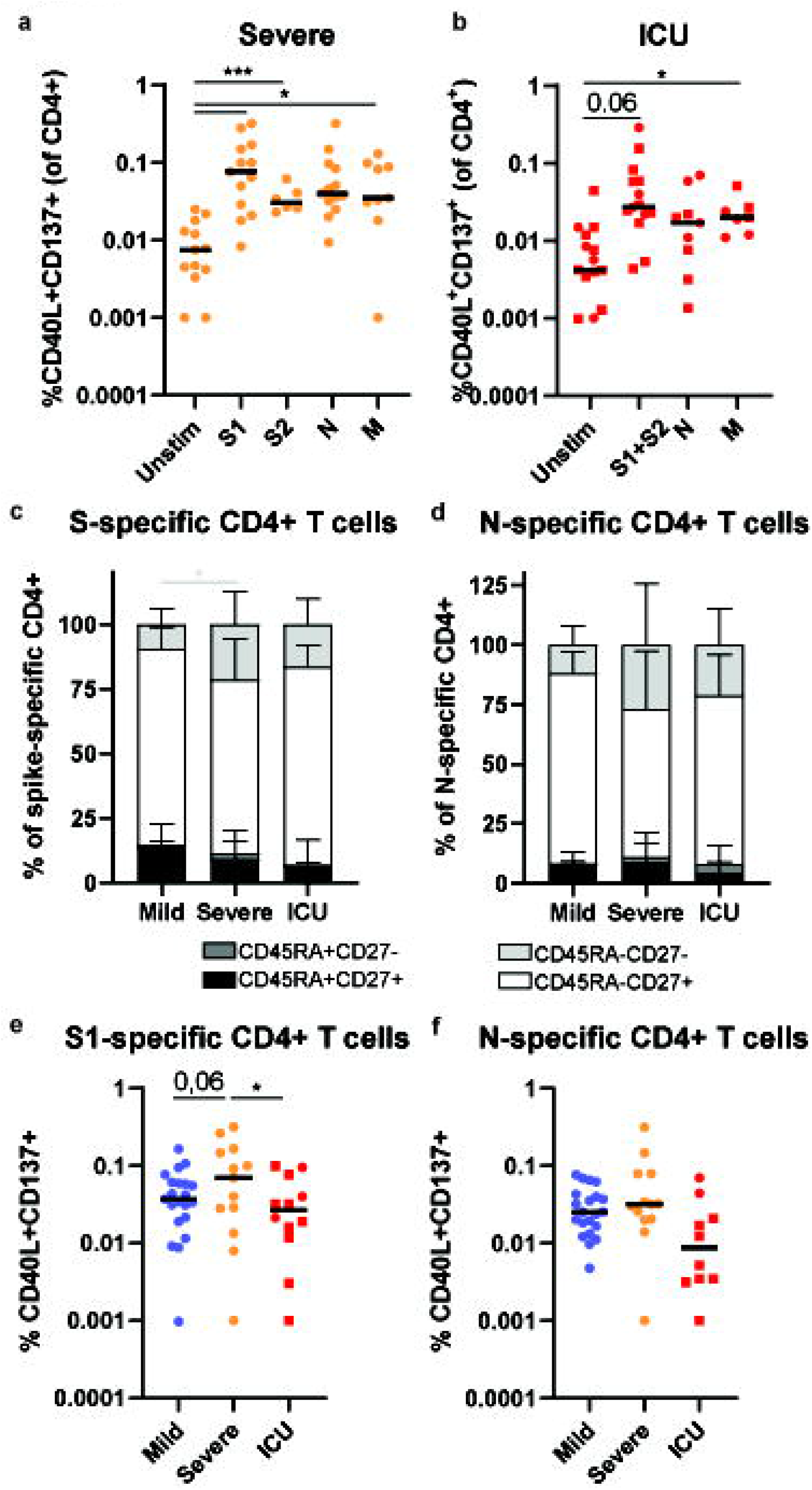
COVID-19 ICU patients have diminished frequencies of SARS-CoV-2 specific CD4+ T cells. **a,b**, Frequencies of S1, S2, or addition of S1 and S2 (S1+S2), N, and M, specific CD4+ T cells were assessed in COVID-19 patients with severe (**a**) or critical (ICU) (**b**) symptoms compared to unstimulated (unstim). Circles indicate samples obtained after clearance of symptoms during recovery phase. Squares indicate samples obtained during ICU stay. **c,d**, The phenotype of the S-(**c**) and N-specific (**d**) CD4+ T cells (CD40L+CD137+) was assessed by measuring CD27 and CD45RA expression to define naïve (black; CD45RA+CD27+), effector (dark grey; CD45RA+CD27-), memory (white; CD45RA-CD27+), and effector memory (light grey; CD45RA-CD27-) cells. **e,f**, The frequencies of S1- (**e**) and N-specific (**f**) CD4+ T cells were compared between the mild, severe and ICU cohorts. Circles indicate samples obtained after clearance of symptoms during recovery phase. Squares indicate samples obtained during ICU stay. **a,b,e,f**, Representative contour plots indicate the gating strategy used. Each symbol indicates a unique sample and the line indicates the median values. Significance of the results was determined using one-way ANOVA with Tuckey’s multiple comparisons test. **c,d,**, Bars show mean values with SD and the significance was determined using two-way ANOVA with Tukey’s multiple comparisons test, color of asterix indicate the groups which are compared. **a**, N=13, N=13, N=13, N=6, N=9 patients for unstim, S1, S2, N, and M conditions respectively. **b**, N=16, N=15, N=9, N=7 patients for unstim, S1+S2, N, and M conditions respectively. **e**, N=21, N=13, N=12 for mild, severe, and ICU patients, respectively. **f**, =21, N=13, N=10 for mild, severe, and ICU patients, respectively.

To determine whether T cell responses were influenced by disease severity, we compared the magnitude of response to S and N between the three cohorts. Overall, by polyclonal aCD3 stimulation, CD4+ T cells from all patient cohorts, in disregard of disease severity, produce relatively similar cytokine profiles. A bias to increased expression of IL-4 and IL-21 production was observed for the by ICU patient cohort, which we attribute to aging effects (Extended Data Fig. 4a). Intriguingly, the severe patient cohort showed higher frequencies of S-specific CD4+ T cells compared to both the mild and ICU cohorts (Fig. 4e). The severe patients also had higher frequencies of N-specific CD4+ T cells (Fig. 4f). While the magnitude of the response to SARS-CoV-2 is important, the quality of the response is also crucial. Therefore, we assessed the cytokine profile of the CD4+ T cells that showed at least a 0.01% response frequency. Regardless of disease severity, SARS-CoV-2-specific CD4+ T cells conformed to a type 1 cytokine profile, predominantly producing IFN-γ and TNF-α (Figure 5a,b, Extended Data Fig. 4b,c). Although ICU patient CD4+ T cells also skewed towards a type 1 response, the S- and N-specific CD4+ T cells produced less IFN-γ compared to the mild group. Although not produced to the same magnitude as TNF-α and IFN-γ, N-specific CD4+ T cells of mild COVID-19 patients also produced significantly more IL-21 than severe or ICU patient CD4+ T cells and a similar trend was observed for S-specific CD4+ T cells. Furthermore, S-specific CD4+ T cells of mild produced significantly more IL-4 than the CD4+ T cells from ICU patients, with a similar trend for the N-specific CD4+ T cells. Thus, the critically ill ICU patients demonstrated a diminished CD4+ T cell response not only in quantity but also in quality of the response.

**Fig. 5:**
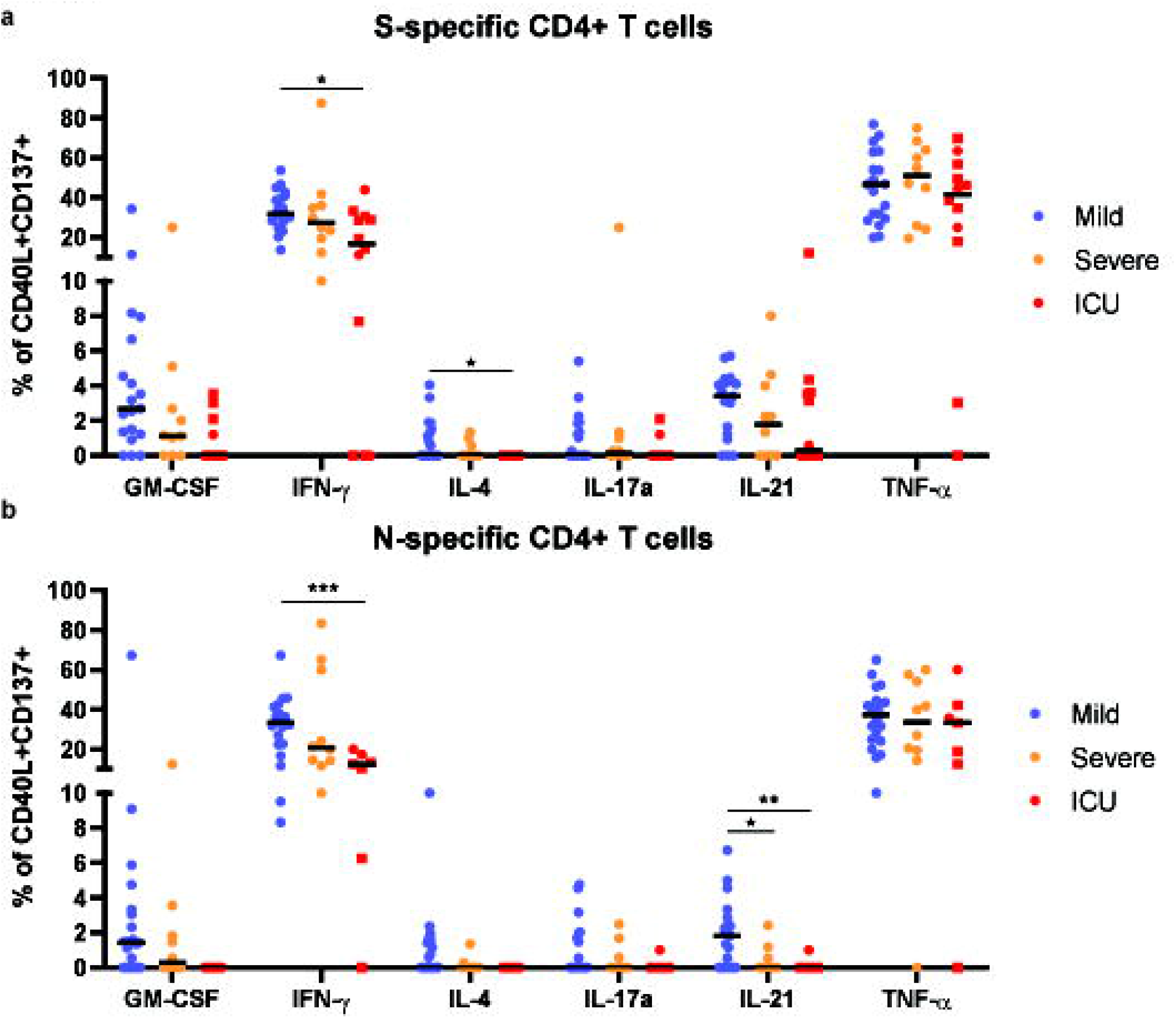
S- and N-specific CD4+ T cells in ICU patients are functionally impaired. **a,b**, Production of GM-CSF, IFN-γ, IL-4, IL-17a, IL-21 and TNF-α production by S- (**a**) and N-specific (**b**) CD4+ T cells (CD40L+CD137+) was determined in mild, severe and ICU patients, in samples that the frequency of S- or N-specific CD4+ T cells (CD40L+CD137+ of CD4+) was at least 0.01%. Each symbol indicates a unique sample and the line indicates the median values. Circles indicate samples obtained after clearance of symptoms during recovery phase. Squares indicate samples obtained during ICU stay. Significance of the results was determined using two-way ANOVA with Tukey’s multiple comparisons test. For S-specific N=19, N=10, and N=12 patients for mild, severe, and ICU cohorts respectively. For N-specific N=21, N=10, and N=7 patients for mild, severe, and ICU cohorts respectively.

### SARS-CoV-2 specific CD4+ T cell responses do not correlate with antibody titers in COVID-19 patients in the ICU

B cell responses in patients with COVID-19 develop from around one week after symptom onset^8^. Coronavirus-neutralizing antibodies primarily target the receptor binding domain (RBD) of the S protein to prevent entry into host cells^24,25^. Neutralizing antibody (nAb) responses begin to develop around week 2 in most patients. IgG responses to the RBD of S was analyzed in the different patient groups. Serum antibody levels correlated with clinical severity (Fig. 6a). Similarly, nucleocapsid (N) IgG titers were increased in the severe and ICU patients compared to the mild group (Fig. 6b). The titers for S-RBD and N correlated well so we focused our analysis on S-RBD from here onwards (Fig. 6c). Since effective antibody responses and isotype switching rely on CD4+ T cell help, we assessed whether SARS-CoV-2 antibody titers were associated with CD4+ T cell responses. As S-specific CD4+ T cell responses were detected in all COVID-19 patients and was indicative for the CD4+ T cell response to the total SARS-CoV-2 antigen pool (Extended Data Fig. 5a), we assessed whether CD4+ T cell responses were associated with antibody titers to SARS-CoV-2 (Fig. 6d). For the mild patient group we found S-specific CD4+ T cell responses, as a proxy for total SARS-CoV-2-specific CD4+ T cell responses, to correlate with the magnitude of anti-S-RBD titers, in line with previous reports^7,26^. No correlation was observed for the severe and ICU patient groups (we acknowledge smaller group sizes) (Fig. 6d, Extended Data Fig. 5c,d). It appeared that SARS-CoV-2 antibody titers developed earlier and stronger in the severe group and especially in the ICU group (Figure 6e, Extended Data Fig. 5e-g), while the S-reactive CD4+ T cell development was highest in the severe group. A significant correlation was observed between the abundance of S-reactive CD4+ T cells and the course of COVID-19 disease for the mild and ICU, but not severe group (Figure 6f, Extended Data Fig. 5h-j).

**Fig. 6:**
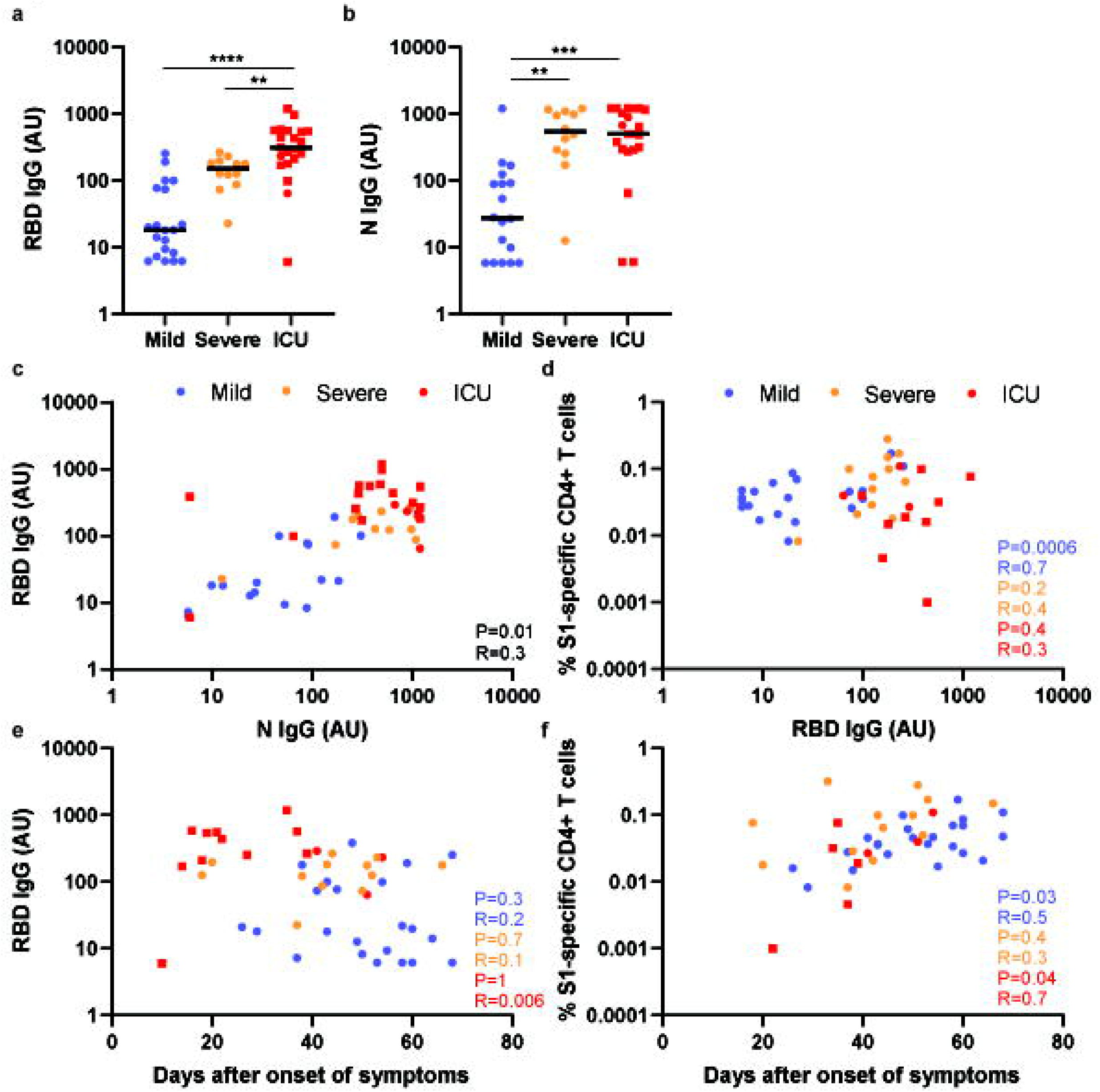
SARS-CoV-2 specific CD4+ T cell responses do not correlate with antibody titers in COVID-19 ICU patients. **a,b**, S-RBD IgG (**a**) and N IgG (**b**) titers were determined in the mild, severe, and ICU cohorts. **c-f**, Correlation between N IgG and RBD IgG titers (**c**), frequency of S-specific CD4+ T cells and RBD IgG titers (**d**), RBD IgG titers and days after symptoms (**e**), and frequency of S-specific CD4+ T cells and days after symptoms (**f**) were assessed in mild, severe, and ICU patients. Circles indicate samples obtained after clearance of symptoms during recovery phase. Squares indicate samples obtained during ICU stay. **c-f**, P and R values are shown for the correlation calculated for each cohort separately. **a-c,e**, N=22, N=12, and N=21 patients for mild, severe, and ICU cohorts respectively. **d,f**, N=22, N=12, and N=21 patients for mild, severe, and ICU cohorts respectively.

## Discussion

In this study we analyzed SARS-CoV-2 specific cellular and humoral responses in COVID-19 patients covering clinical manifestations ranging from mild symptoms to critical illness leading to ICU admission. Both SARS-CoV-2 specific T cell and antibody responses were found in all patients. While we found titers for S-RBD and nucleocapsid, predominant targets for neutralizing antibodies, to correlate with clinical disease severity, the magnitude of SARS-CoV-2-specific T cell responses appeared more variable. Our findings indicate that antibody titers significantly correlated with S-specific CD4+ T cell responses in mild patients that typically recover without special treatment. However, B and T cell responses were imbalanced in the blood of critical ICU patients exhibiting high titers and low virus-specific CD4+ T cell responses. It was not only the magnitude of the CD4+ T cell response that was impaired but also the functionality. SARS-CoV-2-specific CD4+ T cells from ICU patients showed decreased IFN-γ, IL4, and IL-21 production. IFN-γ is not only important for recruitment of other immune cells, but also dampens viral replication^27–30^. Furthermore, IFN-γ and IL-4 can downregulate cell surface expression of ACE2 and inhibit replication of SARS-CoV^31,32^. IL-21 on the other hand can directly act on B cells and regulate germinal center (GC) responses^33^. The increased PD-1 expression and decreased functionality of the SARS-CoV-2 specific CD4+ T cells further strengthen the observation that the anti-viral T cell response is functionally compromised in critical COVID-19 patients.

In line with recent reports, we also detected cross-reactive T cell responses in PBMCs of healthy blood donors obtained before the SARS-CoV-2 pandemic^6,7^. Interestingly, while cross-reactivity with “common cold” coronavirus strains is plausible due to the overlapping epitopes between the viruses, we did not observe significant recognition of 299E and OC43 encoded spike proteins, including the well-conserved S2 domain^34^, in the COVID-19 patients. Nevertheless, the association between the prevalence of cross-reactive T cells and disease severity in patients with COVID-19 needs evaluation in asymptomatic patients including most SARS-CoV-2 infected children^35^. The absence of substantial T cell responses to common-cold coronavirus strains could be causative factor to initiate symptoms, due to delayed control of viral loads.

We found a substantial population of antigen-inexperienced naïve T cells in the BALF of COVID-19 ICU patients. Accompanied with low frequencies of memory T cells with a resident memory (CD69+CD103- or CD69+CD103+) phenotype these data indicate a compromised vascular integrity and epithelial barrier function and thus alveolar leakage. In one ICU patient, in which we were able to obtain representative data of antigen-specific CD4+ T cell in BALF, the S-specific CD4+ T cell response in both BALF and PBMC showed typical phenotypes of circulating cells but not T_RM_, further supporting the notion of vascular leakage. These data support previous findings where increased expansion of T cell clones, indicating possible SARS-CoV-2 specificity, were mainly detected in BALF of moderate but not critical COVID-19 patients^10^. Although these results do not rule out that the lymphopenia in critically ill COVID-19 patients is partly mediated by recruitment of T cells to the lungs, the low number of SARS-CoV-2 specific T cells in the blood and BALF, may rather suggest a compromised expansion or selective depletion, as has been postulated by others^11,36^. Although the role of T_RM_ in controlling SARS-CoV-2 infection remains unexplored, airway virus-specific T cells are elementary in protecting against other respiratory coronaviruses^21^. Future studies analogous to what we have shown for Influenza and RSV should be demonstrated if after clearance of the virus sizable numbers of SARS-CoV-2 specific T cells will remain in the airways.

Our study results imply the requirement of a balanced participation of B and T cells to curb SARS-CoV-2 infection and indicate that T cells cope less efficiently with the rapid viral load peaks observed in COVID-19. SARS-CoV-2 viral load peaks at day 10 after symptom onset while that of previous emerging coronaviruses peaked a few days later^37–40^. Viral loads correlated with disease severity and decreased T cell count, and patient age^40,41^. In line with this, we found patients that developed severe COVID-19, but did not require ICU treatment, to exhibit higher SARS-CoV-2-specific T cell responses alongside high titers for S-RBD and N IgG. On the other hand, critical ICU patients failed to generate substantial T cell responses but generated a strong antibody response already early on. Timing appears to be crucial as the ratio between viral loads and antibody titers during the early phase of disease may be predictive for disease severity^42^. A delayed effective T cell control of viral loads, supplemented with a compromised innate immune function, frequently observed in elderly^43^, could tip the balance towards more severe symptoms mediated by pathological consequences of viral specific antibodies. Such a model may explain the differences observed between the severe and critically ill patient cohorts in our study, where an increased magnitude of anti-S and anti-N specific T cells are associated with lower antibody titers and disease severity. In line with others, we find age to be a risk factor for disease severity. Aging related senescence of potential cross-reactive T cells, combined with a compromised generation of T cell responses to new antigens may make the elderly more susceptible to COVID-19. T cell responses that are generated in critically ill patients may also be dampened by either suppressed expansion or apoptosis induced by a cytokine storm, the main contributor to ARDS observed in critically ill ICU patients^36,44^.

Failure of antiviral T cell responses to control SARS-CoV-2 replication could underlie the hyper-inflammatory responses and associated immunopathology characterizing critically ill COVID-19 patients. These observations seem counterintuitive as B cells require CD4+ T cell help from Tfh for optimal GC responses and class-switching^45^. Circulating Tfh (cTfh) are poorly understood blood counterparts of GC Tfh cells. ICOS+CXCR3+ cTfh correlate with nAb responses upon viral infection or vaccination^46–49^. cTfh increase over the course of SARS-CoV-2 infection^8,15,16^. In our study the cTfh frequency was highest in the ICU patients, patients that also demonstrated the highest antibody titers, with indications of levels decreasing upon recovery. Interestingly CXCR3+cTfh, produce IL-21, which was found less in SARS-CoV-2-specific CD4+ T cells in ICU patients. Although IL-21 is also produced by T_H_2 cells, and serum levels may not be associated with cTfh numbers, we cannot rule out the potential effect of age-related decline in Tfh functions. Although no signs are reported that B cell function is severely compromised in critically ill COVID-19 patients, we encourage investigation of the potential effect of compromised Tfh functions on anti-SARS-CoV-2 reactive antibody seroconversion and affinity maturation. Of note in chronic HCV cTfh show diminished IL-21 secretion but are still capable of supporting memory B cell differentiation and antibody production^50^. Therefore, the role of Tfh in COVID-19 immunity needs further investigation.

On the other hand, B cells also play a role in antigen presentation to activate CD4+ T cells. Interestingly, the quality of the B cell response appears to be specifically altered in critically ill ICU patients with COVID-19. These patients, in contrast to patients with mild symptoms, produce afucosylated IgG antibodies against the S protein^51^. Afucosylated antibodies exhibit increased binding affinity to Fc receptors and enhance antibody dependent cellular cytotoxicity (ADCC), which may account for the hyper immune response in ICU patients. Since crosslinking of Fc receptors is required for optimal antigen-cross presentation and priming of T cells, future studies may direct whether IgG variants without core fucosylation prime T cells equally well as their fucosylated counterparts, and as such contribute to the lower frequency of antigen-specific CD4+ T cells observed in ICU patients. It was recently proposed that signaling through a co-signal receptor may alter the glyco-programming of B cells^51^. Whether CD4+ T cell, in particular Tfh, induced signaling through co-stimulatory molecules or secreted cytokines, such as IL-21, can alter B cell glyco-programming in COVID-19 remains to be elucidated and can be of great importance for vaccine strategies.

Taken together, these data suggest that a balanced participation of B and T cells is required to control SARS-CoV-2 infection and provide rationale for future evaluation of vaccine strategies. Further studies on the direct and indirect interactions between the humoral and cellular adaptive immune response are required for a better understanding of COVID-19 pathophysiology.

## Materials and methods

### Subjects

Blood was collected from ex-COVID-19 patients donating convalescent plasma at Sanquin Blood Bank, Amsterdam, the Netherlands. All donors were diagnosed with SARS-CoV-2 infection by PCR prior to inclusion. All patients admitted to the intensive care unit (ICU) of the VU medical center that received a diagnostic BAL were included in this study. Sanquin Blood Supply Foundation, Amsterdam, the Netherlands supplied the buffy coats of regular donors used as unexposed healthy controls.

### Ethics approval

Data and samples were collected only from voluntary, non-remunerated, adult donors who provided written informed consent as part of routine donor selection and blood collection procedures, that were approved by the Ethics Advisory Council of Sanquin Blood Supply Foundation. Data and samples from ICU patients of the Amsterdam UMC, location VUmcare part of the Amsterdam UMC COVID-19 biobank (2020-182). The Amsterdam UMC COVID-19 biobank stores leftover patients samples that were collected for diagnostics purposes, which is approved by Review Committee Biobank of the Amsterdam UMC (2020-065). The study is in accordance with the declaration of Helsinki and according to Dutch regulations

### Antibody titers

Antibody titers were determined by ELISA analogously. Briefly, samples were tested at 100 – 1200 fold dilutions (in PBS supplemented with 0.1% polysorbate-20 and 0.3% gelatin (PTG) in microtiter plates coated with RBD or (nucleocapsid) NP and incubated for 1h at RT. Both proteins were produced as described^51^. After washing, 0.5 μg/mL HRP-conjugated antihuman IgG (MH16, Sanquin) was added in PTG and incubated for 1h. Following enzymatic conversion of TMB substrate, absorbance was measured at 450 nm and 540 nm and the difference used to evaluate antibody binding by comparison to a reference plasma pool of convalescent COVID-19 patients.

### Isolation of peripheral blood mononuclear cells and BALF mononuclear cells

Peripheral blood mononuclear cells (PBMCs) were isolated from heparinized blood samples using standard Ficoll-Paque density gradient centrifugation. During diagnostic bronchoscopy lungs were flushed with 40ml NaCl to collect bronchoalveolar lavage fluid (BALF). BALF was centrifuged (to collect BALF supernatant; 300g, 10min) and the cell pellet was resuspended in 2mM dithiotreitol (Sigma, Zwijndrecht, the Netherlands). After 30 min at 4°C, cells were washed with PBS+1%BSA and mononuclear cells were isolated using Ficoll-Paque density gradient centrifugation. PBMC and BALF MC samples were either used directly for experimentation or cryopreserved in liquid nitrogen until further analysis.

### SARS-CoV-2 peptide pools

The details of the peptide pools (JPT, Germany) used in this study are listed in Extended Data Table 2. Peptide pools against spike (S1 and S2), nucleocapsid (N), membrane protein (M), envelope protein (E), ORF3A and pool of open reading frame (ORF), non-structural (NS) and uncharacterized proteins referred to as ORF pool; containing NS6, NS7B, NS7A, NS8, ORD10, ORF9B and Y14, were used when indicated. When samples allowed, the S1 and S2 were added up and referred to as S1+S2 in the figures. Peptide pools were used at a final concentration 100 ng/mL.

### In vitro stimulation assay

Cryopreserved PBMC and BALF samples were thawed. Alternatively, fresh PBMC samples were used directly after isolation. For stimulation experiments with BALF, paired PBMC samples were first labelled with cell trace violet (CTV, Thermofisher) according to manufacturer’s instructions. Thereafter, both CTV labeled PBMCs and BALF samples were stained with CD69 BUV395. For BALF experiments, paired CTV labelled PBMCs were mixed with the BALF MCs at a ratio of 1:1. For stimulation experiments with only PBMCs, approximately 2 million cells were used per condition. Cells were incubated with peptide pools at 37°C (5% CO2). Unstimulated cells with no added peptides was used a negative control. Activation with soluble αCD3 (Thermofisher, clone HIT3A) was used as a positive control. After two hours, Brefeldin A (Thermofisher) was added and the cells were further incubated at 37°C overnight.

### Flow cytometric analysis

PBMCs and BALF MCs were stained with combinations of the following antibodies: anti-CD3 BUV661 (BD, clone UCHT1), anti-CD4 BUV737 (BD, clone SK3), anti-CD8 BUV805 (BD, clone SK1), anti-CD45RA BUV563 (BD, clone HI100), anti-CD27 BV650 (Biolegend, clone O323), anti-CD69 BUV395 (BD, clone FN50), anti-CD103 BV711 (Biolegend, clone Ber-ACT8), anti-PD-1 BB700 (BD, clone EH12.1), anti-CXCR5 AF488 (BD, clone RF8B2), anti-ICOS BV711 (Biolegend, clone C398.4A), anti-CD40L PeDazzle594 (Biolegend, clone 24-31), anti-CD137 PeCy7 (Thermofisher Scientific, clone 4B4-1), anti-TNF BV510 (Biolegend, clone Mab11), anti-IFNγ BV785 (Biolegend, clone 4S.B3), anti-IL17a BV605 (Biolegend, clone BL168), anti-IL-4 BV421 (Biolegend, clone MP4-25D2), and anti-IL-21 eFluor660 (Thermofisher, clone 3A3-N2).

Cells were stained according to the manufacturer’s instructions and analyzed in PBS 0.5% FCS. For intracellular staining, cells were fixed and permeabilized with the Foxp3 Staining Buffer Set (Thermofisher). Samples were acquired on a BD FACSymphony. Data analysis was performed using FlowJo (TreeStar, Version 10.0.7).

### Statistics

One- and two-way ANOVA and Tukey’s multiple comparisons test using GraphPad Prism 8 were used to determine the significance of our results as indicated in figure legends. p value of less than 0.05 was considered statistically significant (* p<0.05; ** p<0.01; *** p<0.001; **** p<0.0001). Pearson’s correlations were calculated to define correlations throughout the manuscript.

## Acknowledgements

We would like to thank all patients and donors who participated in this study. Furthermore, we’d like to thank the people of the Sanquin COVID-19 biobank and the ARTDECO study consortium for the collection and processing of samples. We would also like to extend our gratitude to Erik Mul, Simon Tol, and Mark Hoogenboezem of the Sanquin Central Facility for their assistance. We would also like to thank Gerard van Mierlo and Gestur Vidarsson for their contribution in the study. This study was funded by Sanquin Blood Supply project grant PPOC, Jochem van Loghem and the Dutch Research Counsil Veni grant.

## Author contribution

AO, AtB, PH designed the study. EN, LH, and AS contributed to the collection and processing of the ICU patient BALF and PBMC samples. ES, FS, HV, and EvdS contributed to setting up the Sanquin COVID-19 Biobank. AO, CG, NK, and PH processed samples of the Sanquin COVID-19 Biobank. SC contributed to the collection of clinical data. AO, AS, CG, NK, PH, TR, and BH performed experiments in this manuscript. AO and TR performed the data analysis. AO, AtB, RvL, and PH wrote the manuscript. All authors approved the manuscript.

