## Supplementary figures and images for "Divergent SARS-CoV-2-specific T and B cell responses in severe but not mild COVID-19"

### Supplementary figures_rPH-01.tif

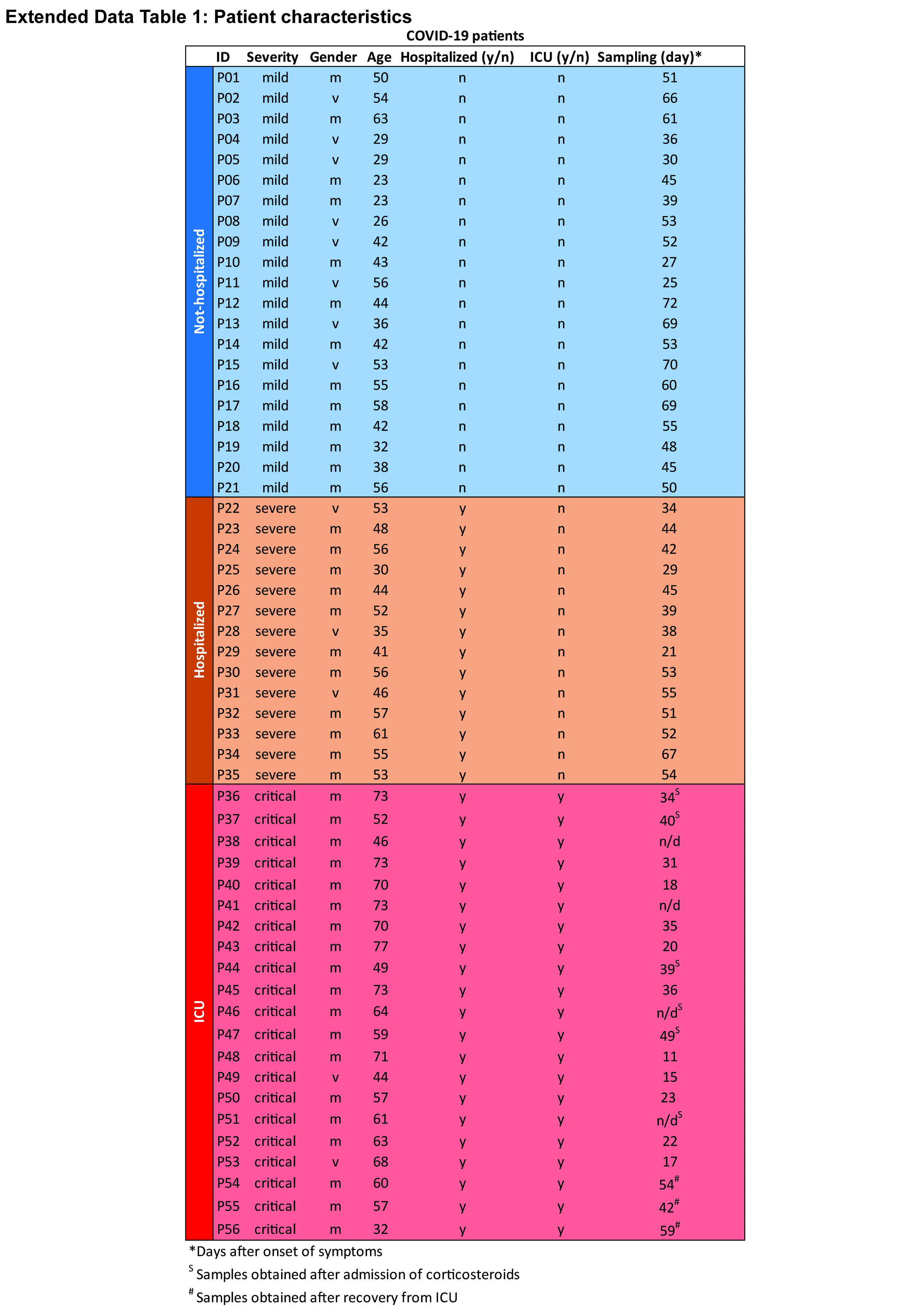

### Supplementary figures_rPH-02.tif

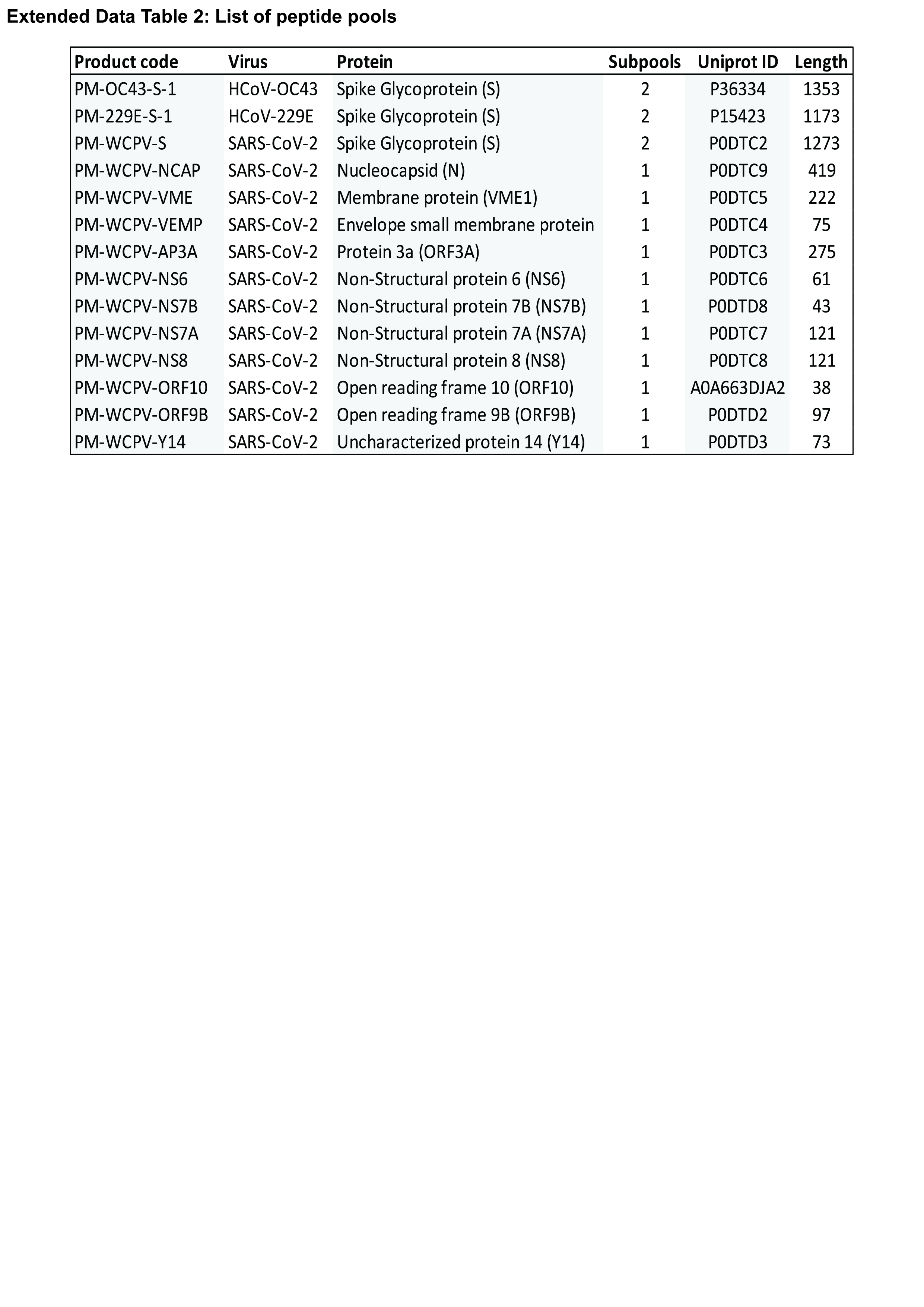

### Supplementary figures_rPH-03.tif

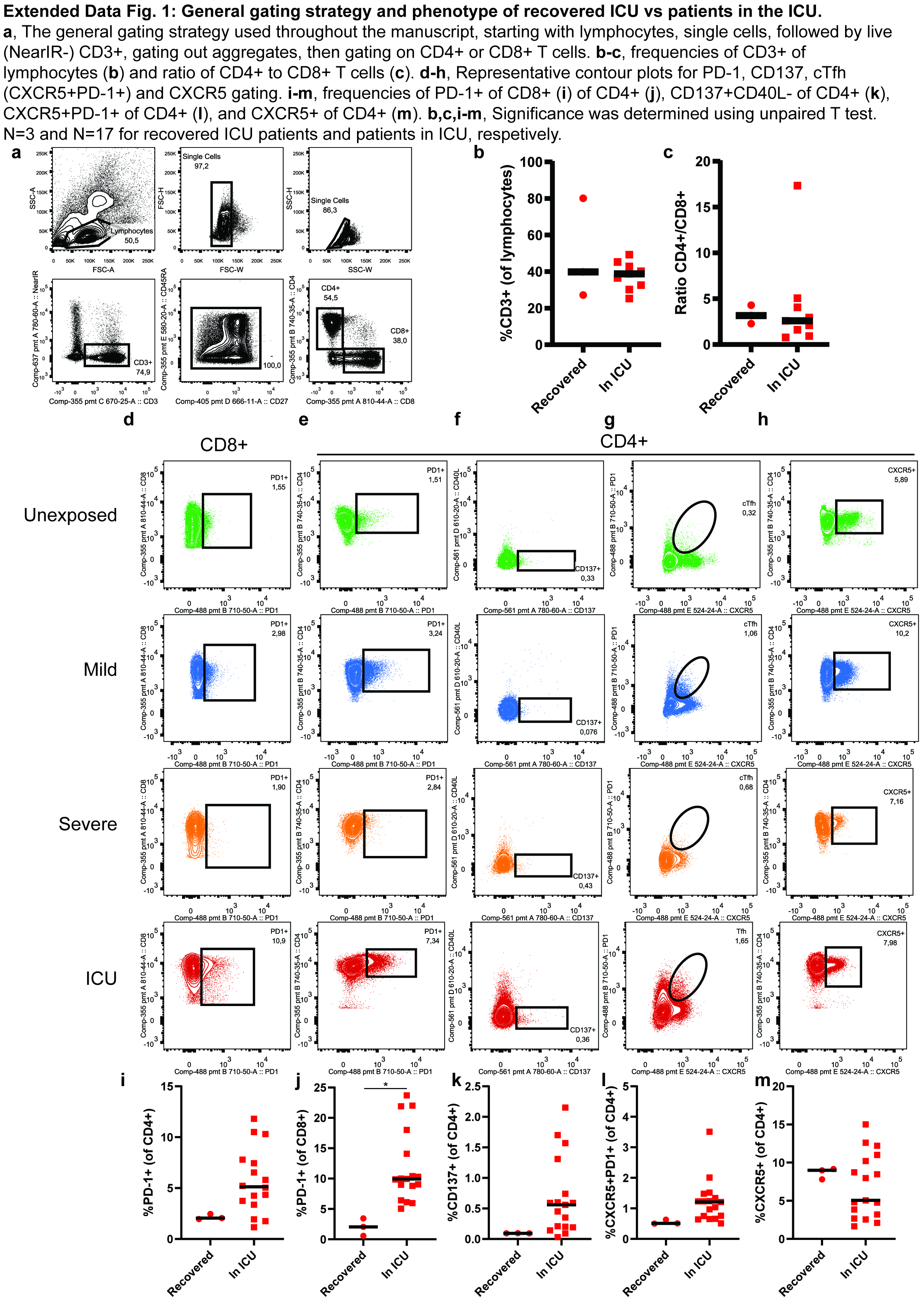

### Supplementary figures_rPH-04.tif

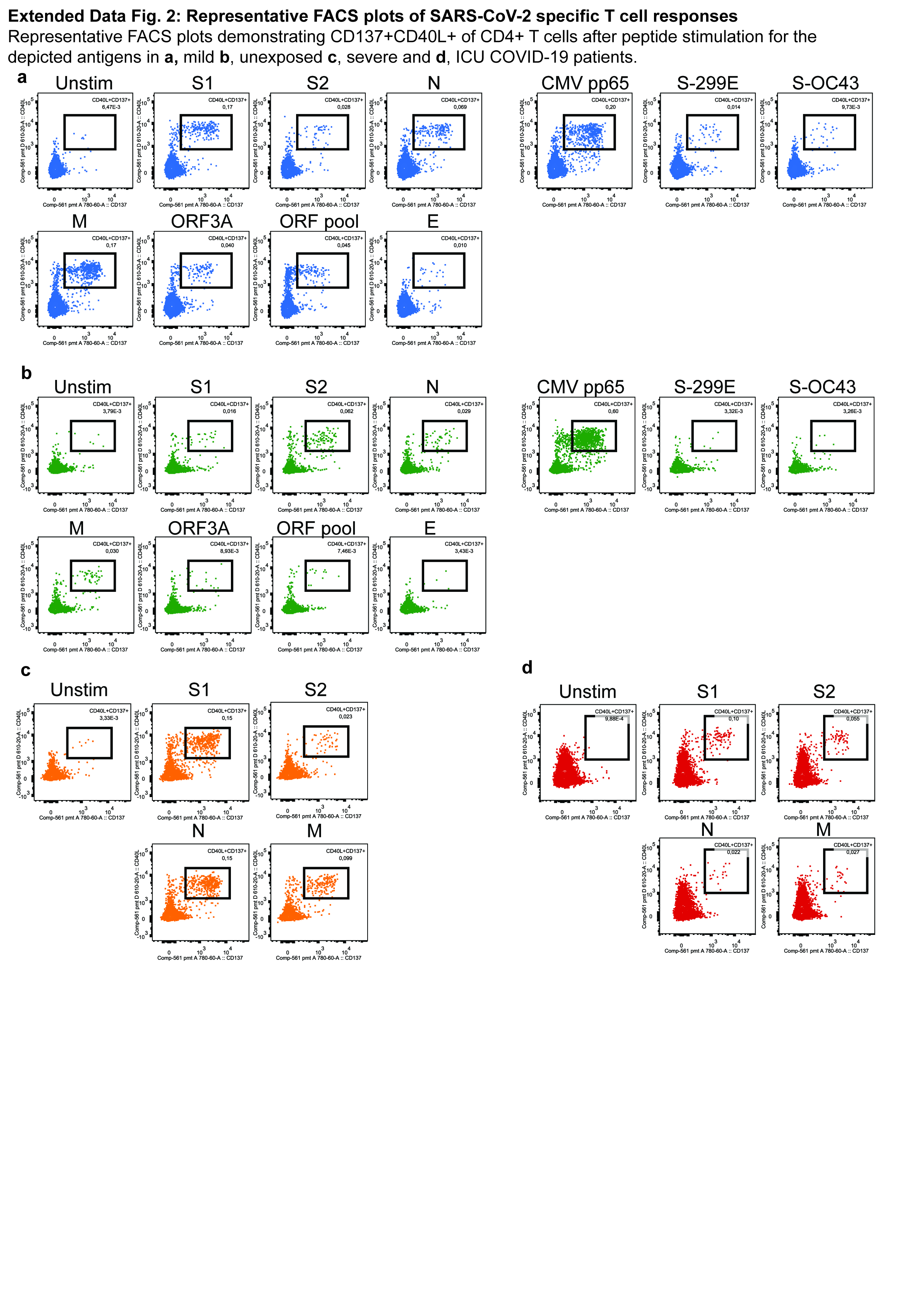

### Supplementary figures_rPH-05.tif

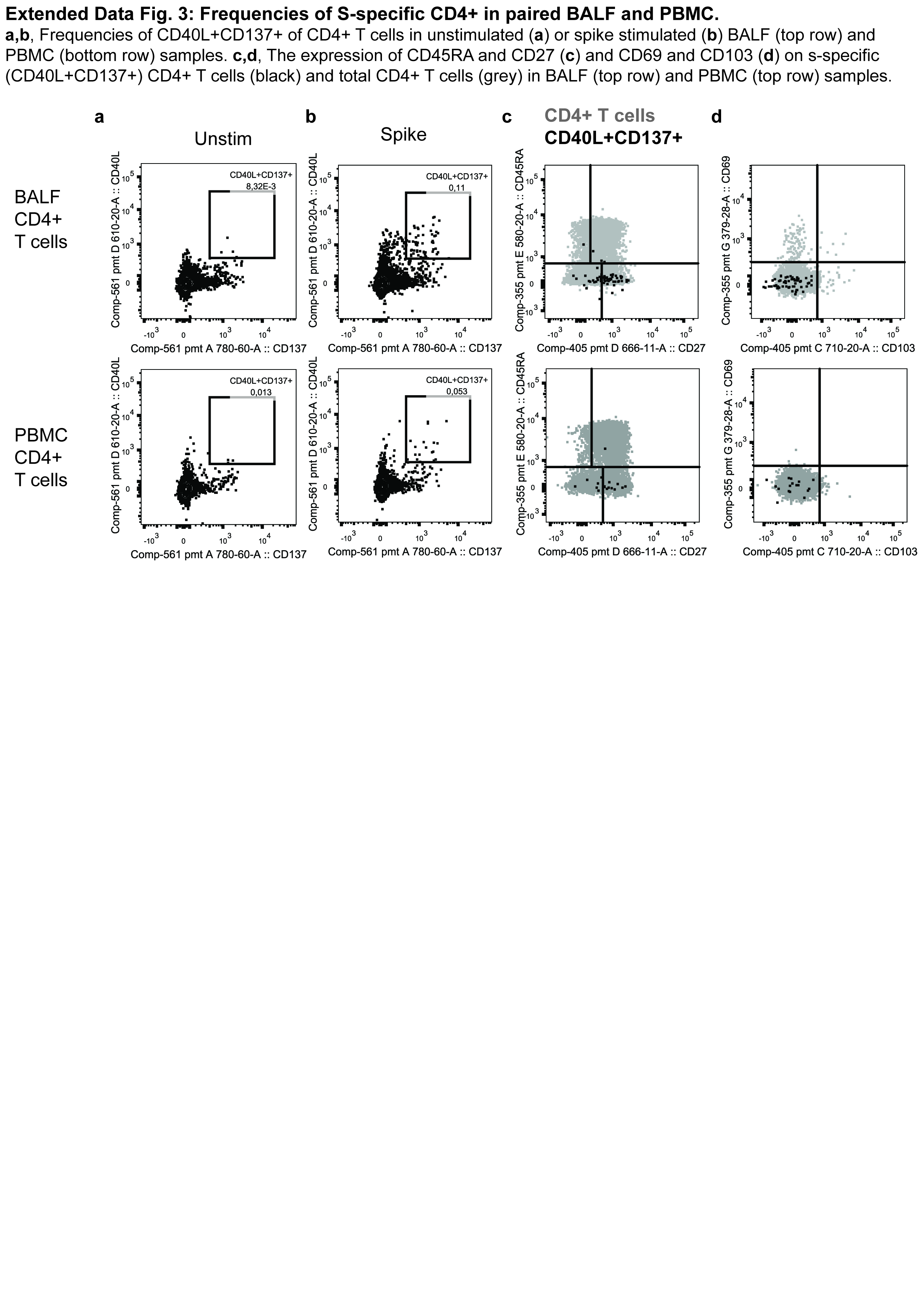

### Supplementary figures_rPH-06.tif

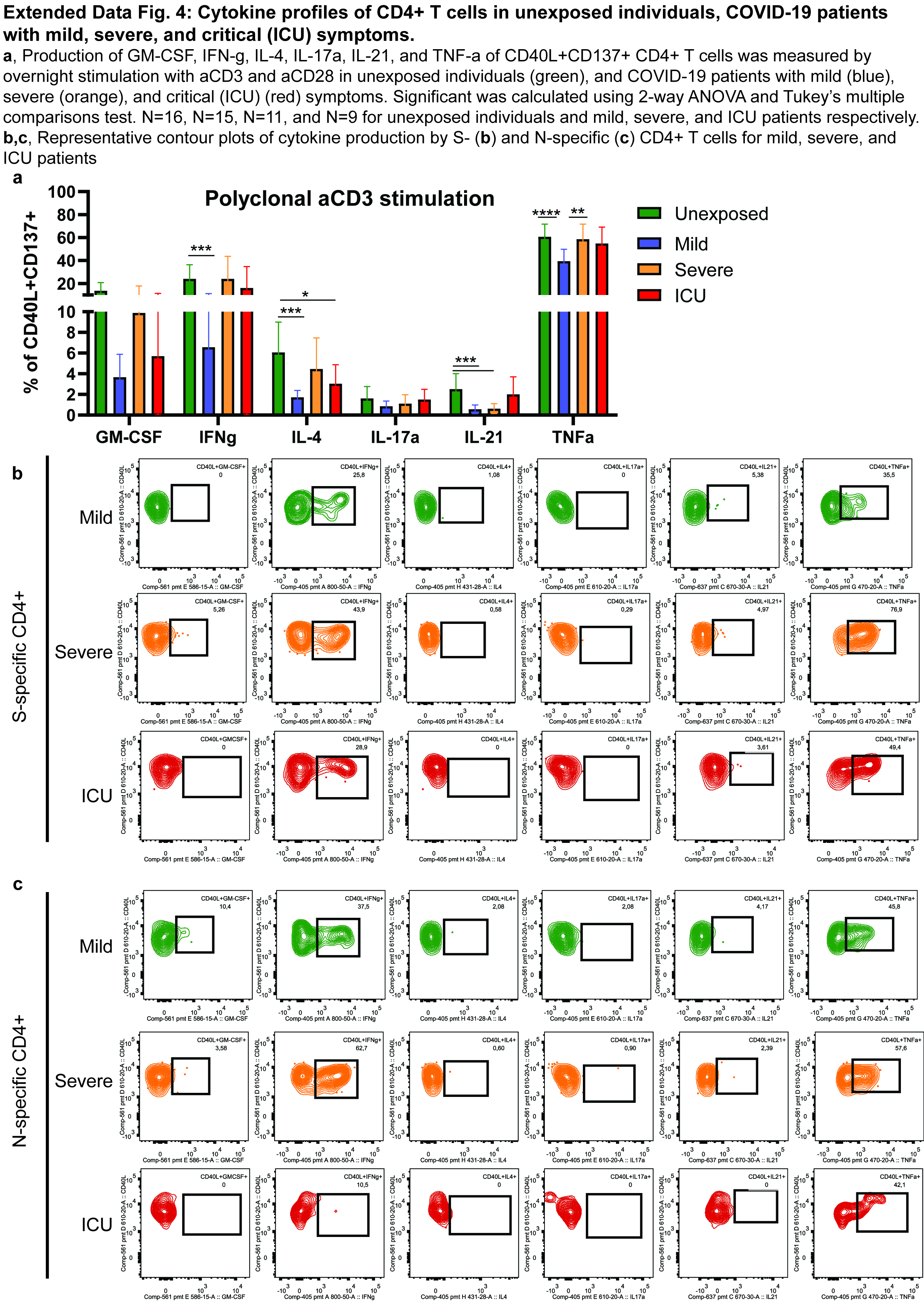

### Supplementary figures_rPH-07.tif

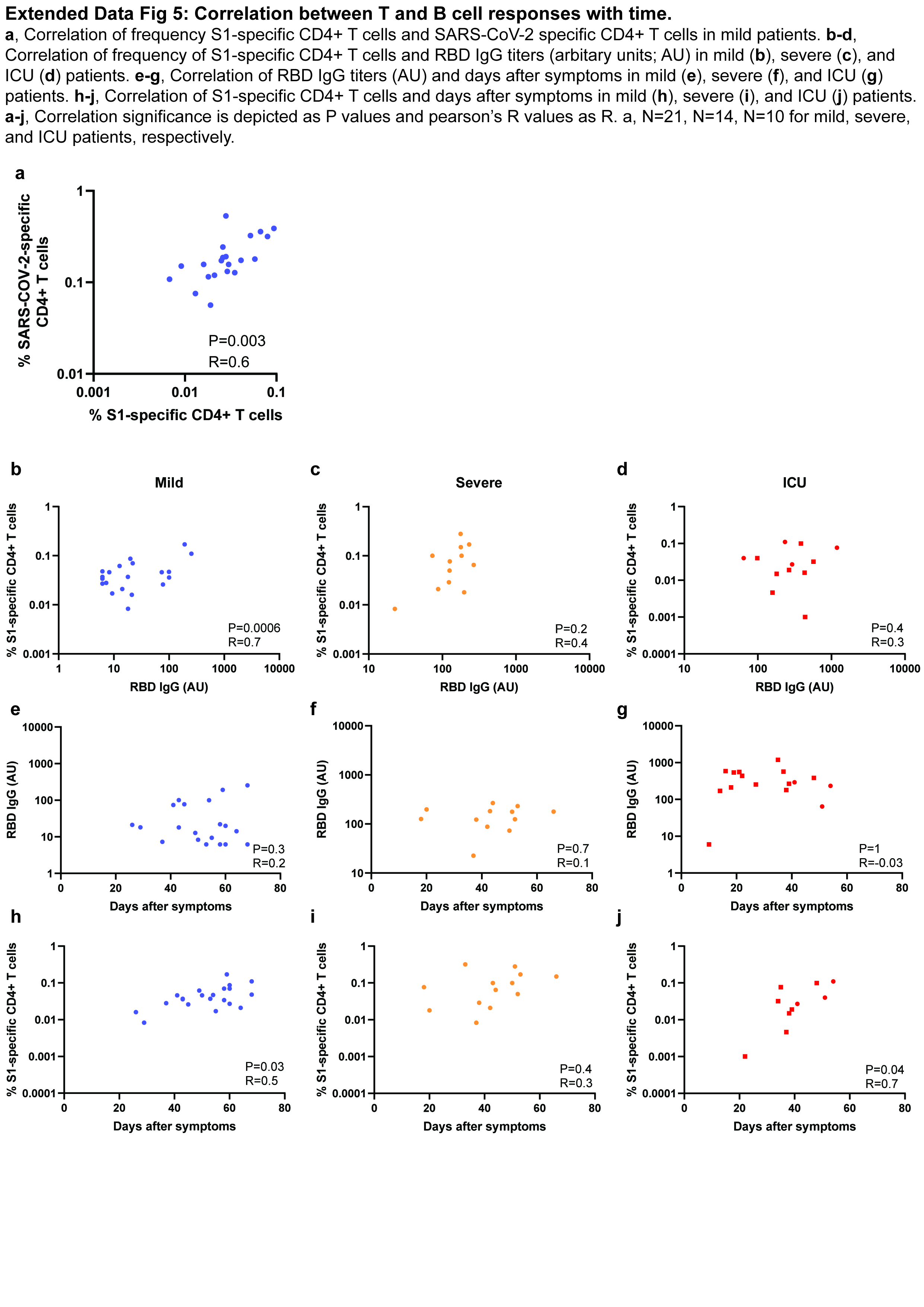
